# Time-lapse Imaging of Microglial Activity and Astrocytic Calcium Signaling Reveals a Neuroprotective Effect of Cannabidiol in the Subacute Phase of Stroke

**DOI:** 10.1101/2022.05.31.494189

**Authors:** Erika Meyer, Phillip Rieder, Davide Gobbo, Gabriela Cândido, Anja Scheller, Rúbia Maria Weffort de Oliveira, Frank Kirchhoff

## Abstract

Pharmacological agents that limit secondary tissue loss and/or improve functional outcomes after stroke are still limited. Cannabidiol, the major non-psychoactive component of *Cannabis sativa*, has been proposed as a neuroprotective agent against experimental focal cerebral ischemia. The effects of cannabidiol have generally been related to the modulation of neuroinflammation, including the control of glial activation and the toxicity exerted by pro-inflammatory mediators. However, so far, most information concerning cannabidiol neuroprotective effects was obtained from histological and biochemical *post-mortem* assays. To test whether the effects of cannabidiol on glial cells could be also detected *in vivo*, we performed time-lapse imaging of microglial activity and astrocytic calcium signaling in the subacute phase of stroke using two-photon laser-scanning microscopy. First, C57BL/6N wild-type mice underwent either sham or transient middle cerebral artery occlusion surgery. The animals received intraperitoneal injection of vehicle or cannabidiol (10 mg/kg) 30 min, 24 h, and 48 h after surgery. One day later the neurological score test was performed. Brain tissue was processed to evaluate the neuronal loss and microglial activation. Transgenic mice with microglial expression of the enhanced green fluorescent protein and astrocyte-specific expression of the calcium sensor GCaMP3 were used to access *in vivo* microglial activity and astrocytic calcium signaling, respectively. The animals were submitted to the same experimental design described above and to imaging sessions before, 30 min, 24 h and, 48 h after surgery. Astrocytic calcium signaling was also assessed in acutely isolated slices 5 h after transient middle cerebral artery occlusion surgery in the presence of perfusion or cannabidiol solution. Cannabidiol prevented ischemia-induced neurological impairments as well as protected against neuronal loss in ischemic mice. Cannabidiol also reduced ischemia-induced microglial activation, as demonstrated in fixed tissue as well in *in vivo* conditions. No difference in the amplitude and duration of astrocytic calcium signals was detected before and after the middle cerebral artery occlusion *in vivo*. Similarly, no significant difference was found in the astrocytic calcium signals between contra and ipsilateral side of acutely isolated brain slices. The present results suggest that the neuroprotective effects of cannabidiol after stroke may occur in the subacute phase of ischemia and reinforce the strong anti-inflammatory property of this compound.

## INTRODUCTION

Stroke is one of the most important causes of morbidity and mortality worldwide. Globally, from 1990 to 2019, the number of incident strokes and deaths due to stroke increased by 70% and 43% (GBD, 2021). Stroke survivors are particularly vulnerable to the stroke-secondary consequences that include sensory, motor and cognitive impairments, as well as mood dysfunction. Despite some recent advances in ischemic stroke caused by large-vessel occlusions, such as mechanical thrombectomy, a neuroprotective pharmacological agent that reduces the consequences of stroke remains needed in clinical practice.

Resident microglia are the first cell of the brain to respond to the pathophysiological changes induced by an ischemic stroke. Several studies have demonstrated the involvement of microglia in cerebral ischemic conditions, but their contribution to the progression of stroke remains under debate, despite the research effort in this field (Kitamura et al., 2004, 2005; Lalancette-Hébert et al., 2007; Jolivel et al., 2015; Szalay et al., 2016). During the acute phase of cerebral ischemia (CI), astrocytes undergo also important morphological modifications, such as hyperplasia and hypertrophy, followed by the production of neurotoxic substances, possibly aggravating the ischemic lesion (Buskila et al., 2005; Basic Kes et al., 2008).

The complex pathophysiology of ischemic stroke may be the reason for the ineffectiveness of treatments that act only on some mechanism of the ischemic cascade. Thus, therapies acting on multiple pathophysiological processes can offer promising results in stroke research. Due to its pleiotropic action and good safety profile cannabidiol (CBD) has been emerging as a candidate for the multifactorial treatment of stroke and its consequences.

Current evidence for the neuroprotective effect of CBD in stroke is predicated on preclinical settings. CBD decreased infarct size, increased cerebral blood flow and improved neurological scores and motor coordination in a model of stroke induced by middle cerebral artery occlusion (MCAO) in mice (Hayakawa et al., 2004; Mishima et al., 2005; Hayakawa et al., 2008). CBD also protected against neuronal death and reduced microglial activation resulting in a decrease of the infarct size in MCAO mice (Hayakawa et al., 2007; Hayakawa et al., 2008; Ceprián et al., 2017). A reduction of microglia and astrocytic reactivity were observed in mice after bilateral common carotid artery occlusion (Schiavon et al., 2014; Mori et al., 2017), a model of transient global CI. Although these studies indicate an effect of CBD in decreasing microglial and astrocytic activation after an ischemic insult, so far these observations were only obtained from immunohistochemical assays.

In the present study we were particularly interested in utilizing *in vivo* imaging to examine how microglia behave after CBD treatment during the early phase of CI induced by MCAO. Using genetically encoded Ca^2+^ indicator and two-photon imaging we also asked if CBD treatment would impact astrocytic Ca^2+^ signaling in the penumbra area of ischemic mice.

## MATERIALS AND METHODS

### Ethics statement

All animal procedures were carried out at the University of Saarland in strict accordance with the recommendations to European and German guidelines for the welfare of experimental animals and approved by the Saarland state’s “Landesamt für Gesundheit und Verbraucherschutz” in Saarbrücken/Germany (animal license number: 36/2016).

### Animals

Mice were housed at the animal facility of the Center for Integrative Physiology and Molecular Medicine (CIPMM) in Homburg under conditions of controlled temperature (22 ± 1 °C) and a 12 h light-dark cycle, with food and water *ad libitum.* Experiments were conducted with 12- to 15-week-old male and female C57BL/6N wild-type (WT) and transgenic mice. To image microglia *in vivo*, the knock-in mouse line TgH(CX_3_CR_1_-EGFP) was used. To visualize astrocytic calcium signals, inducible CreERT2 DNA recombinase knock-in mice TgH(GLAST-CreERT2) were crossed to the floxed reporter mice TgH(Rosa26-CAG-fl-stop-fl-GCaMP3-WPRE). To simplify, in this study the mouse lines are termed CXCR^EGFP^ and GLAST^GCAMP3^, respectively. Further mouse line information and respective constructs are listed below (Table 1).

**Table 1.**
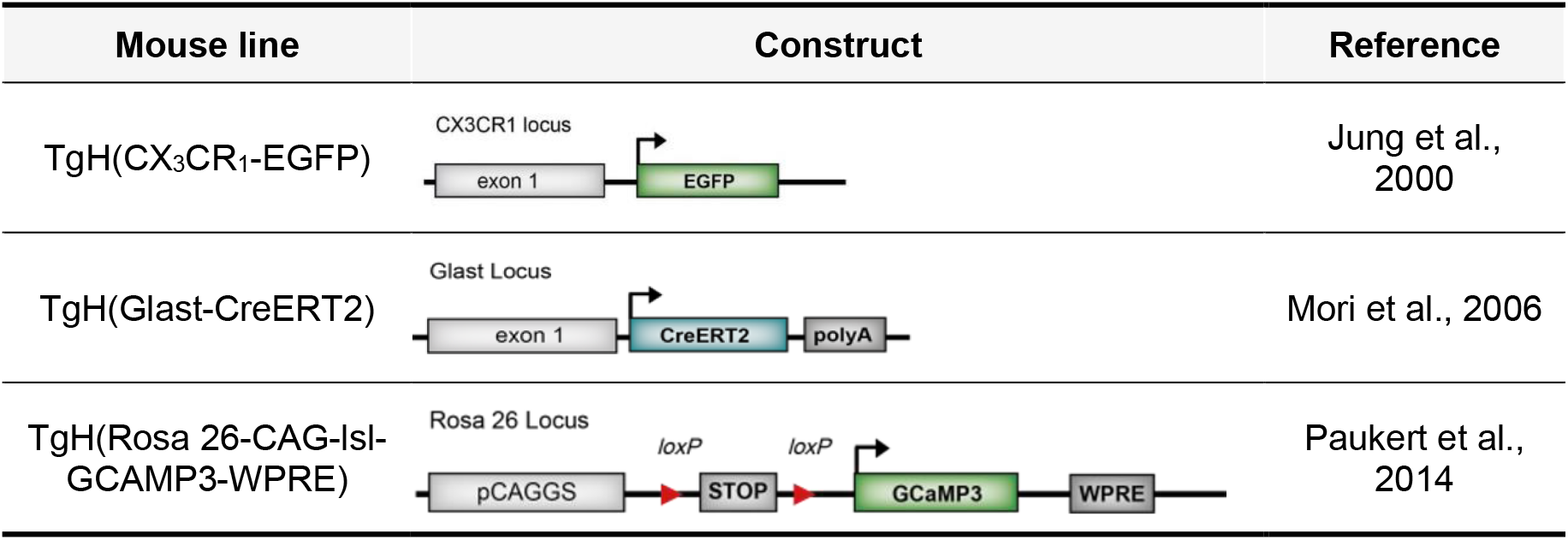
Mouse constructs.

### TgH (CX_3_CR_1_-EGFP)

The mouse line was maintained in the C57BL/6N background and only heterozygous mice were used. In this mouse line, microglia express the enhanced green fluorescent protein (EGFP). EGFP expression is achieved through placement of the EGFP reporter gene into the *Cx3cr1* locus encoding the fractalkine receptor CX3CR1 (Jung et al., 2000).

### TgH (Glast-CreERT2)

In this transgenic mouse line, CreERT2 is knocked into the locus of the L-Glutamate/L-aspartate transporter (GLAST; Mori et al., 2006). The Cre DNA recombinase is bound to the ligand-binding domain of the modified estrogen receptor (ERT2) and remains in the cytoplasm. After tamoxifen administration, this fusion protein translocates into the nucleus and mediates recombination. For this study, only heterozygous animals were used.

### TgH (Rosa 26-CAG-fl-stop-fl-GCaMP3-WPRE)

GCaMP3 is a Ca^2+^ indicator, consisting of the M13 fragment from myosin light chain kinase (M13), EGFP, and calmodulin. The M13 domain is bound at the N-terminus of the EGFP and is target sequence for calmodulin. After binding of Ca^2+^ to calmodulin, the conformation of the protein is changed and the fluorescence intensity of EGFP is thereby enhanced (Nakai et al., 2001). The expression of GCaMP3 is controlled by the CAGGS promoter (Niwa et al., 1991) and the construct is inserted into the Rosa26 locus. Upon tamoxifen administration, the CreERT2 recombinase leads to the deletion of a floxed STOP cassette ahead of the GCaMP3 sequence allowing the inducible expression. The WPRE (woodchuck hepatitis virus posttranscriptional regulatory element) leads to the early exit of mRNA from the nucleus and enhances mRNA stability (Paukert et al., 2014). For this study, only homozygous animals were used.

### Genotyping

Genomic DNA extraction was performed on tail biopsy or ear punch samples. Tissue digestion for DNA extraction and subsequent polymerase chain reaction (PCR) was performed with the prepared sample buffer from REDextract-N-Amp PCR KIT (Sigma-Aldrich, St. Louis, United States) or DreamTaq™ Hot Start Green DNA Polymerase (Thermo Fischer Scientific, Walthan, United States) and different oligonucleotide primers (Table 2). First, the samples were incubated with 62.5 μl of extraction solution for 10 min by shaking. Next, the solution was incubated for 20 min at 95 °C. Following cooling down to RT, 50 μl of neutralization buffer was given to the samples. The reactions were run in 96-well PCR plates in Thermocyclers (PEQLAB Biotechnologie GmbH, Erlangen, Germany). Gel electrophoresis was run on 1.5 - 2 % agarose gels with ethidium bromide (*f.c*. 0.015 %) and, lastly, exposed and documented with the Quantum Gel documentation system (PEQLAB Biotechnologie GmbH, Erlangen, Germany).

**Table 2.**
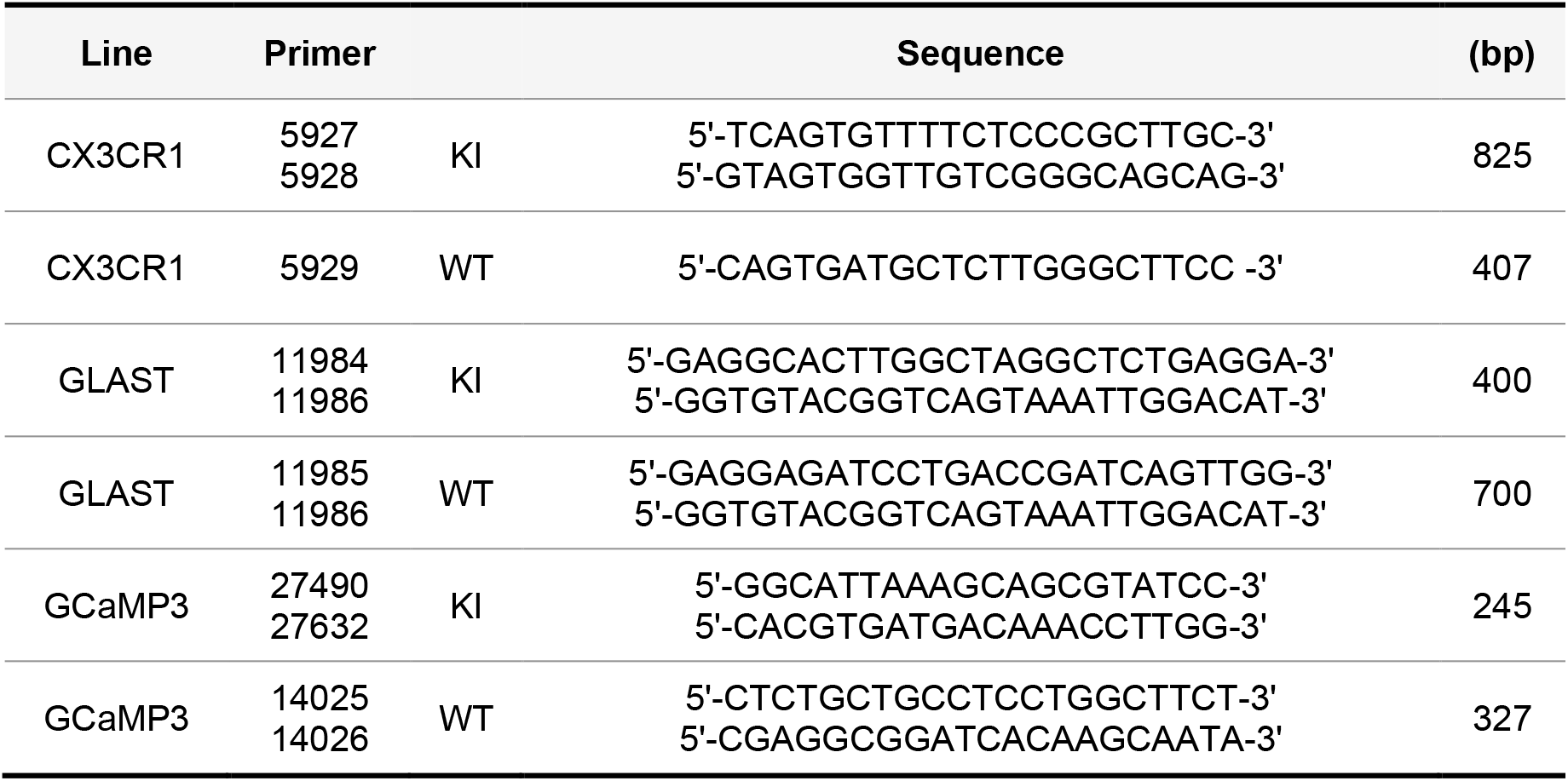
Genotyping primers.

### Drugs

#### Tamoxifen induced gene recombination

Tamoxifen (Sigma-Aldrich, St. Louis, United States) was dissolved in Miglyol (Sigma-Aldrich, St. Louis, United States) at a concentration of 10 mg/ml, aliquoted and stored at 4 °C. The mice received injections of tamoxifen (100 mg/kg, *i.p*.), once per day, for five consecutive days (Jahn et al., 2018). Animals were treated at 7 weeks of age, resulting in 5 weeks intervals before the experiment start (Fig. 1C).

**Figure 1.**
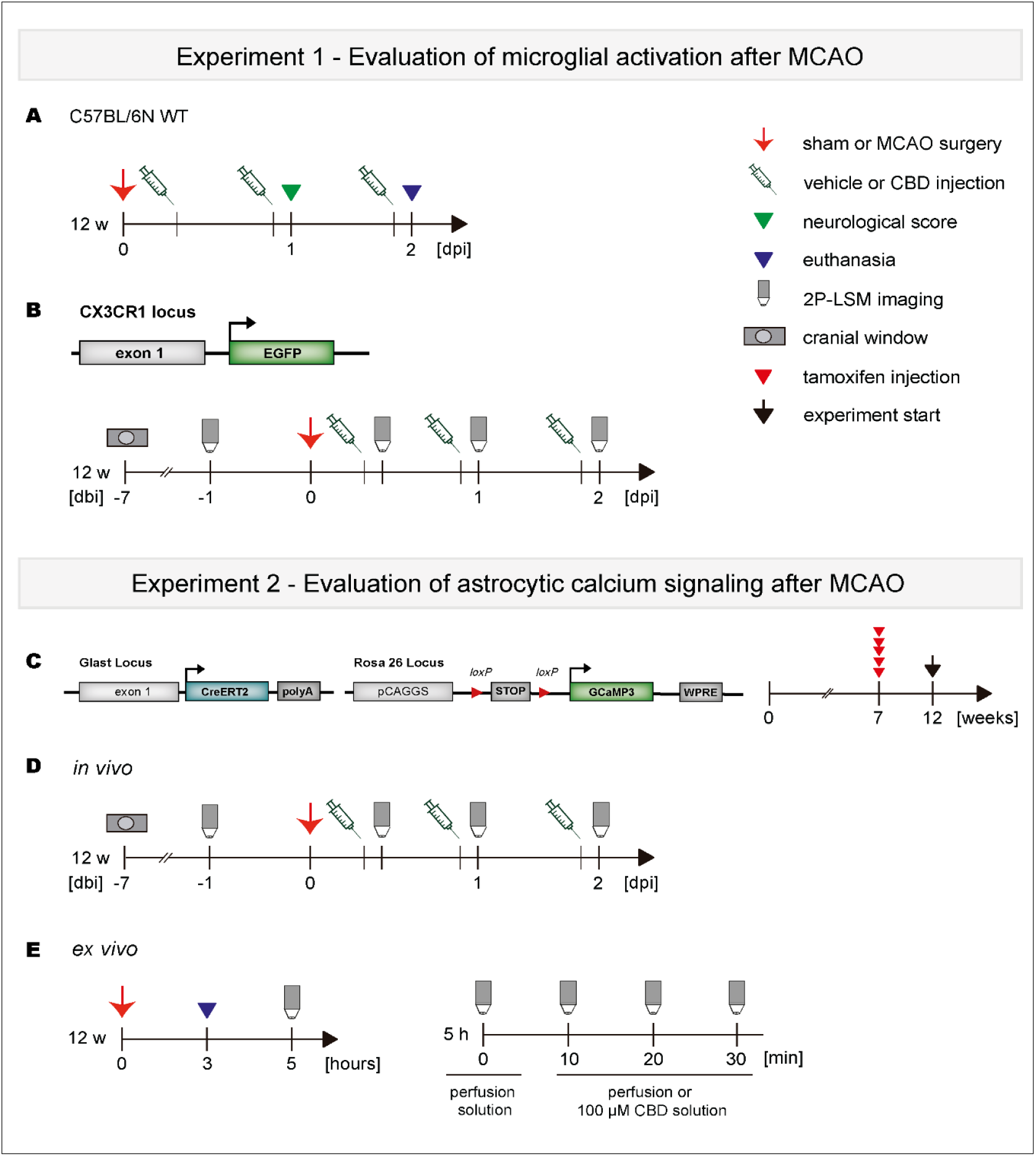
Experimental design. **A-B** In Experiment 1, C57BL/6N WT mice were subjected to sham or MCAO surgery. Vehicle or 10 mg/kg CBD was administered (*i.p.*) 30 min, 24, and 48 h after reperfusion. One day after reperfusion the neurological score was carried out. One day after behavioral testing the animals were euthanized and their brains were collected for analysis. In another protocol, CXCR^EGFP^ mice underwent cortical craniotomy. After one week of recovering, the mice underwent the MCAO surgery. The MCAO mice were treated with vehicle or 10 mg/kg CBD following the same administration protocol cited above. 2P-LSM imaging was performed before MCAO (baseline), 30 min, 24, and 48 h after reperfusion. **C** In experiment 2, the tamoxifen-inducible GLAST-CreERT2 x GCaMP3 mouse line was employed for expression of the Ca^2+^ sensor GCaMP3 in astrocytes. GCaMP3 expression in astrocytes was induced five weeks prior to the experimental start. **D** Mice underwent, first, the cortical craniotomy, and six days later, the MCAO surgery. The mice were submitted to the same treatment- and *in vivo* imaging protocol as described in B. **E** For the *ex vivo* imaging procedure, the mice underwent MCAO, and 3 h later were euthanized and had their brains processed. The brain slices (contra- and ipsilateral site) were incubated in perfusion- or 100 μM CBD solution and imaged over time. CBD: cannabidiol; MCAO: middle cerebral artery occlusion.

#### Cannabidiol treatment

Cannabidiol (CBD; THC Pharma, Frankfurt, Germany) was diluted in 1 % Tween 80 in sterile isotonic saline (vehicle). The animals were randomly assigned to receive injections (*i.p.*) of CBD 10 mg/kg or vehicle 30 min, 24, and 48 h after surgery (Fig. 1A, B, D).

#### Surgeries

For all described surgeries, the procedures were performed under conditions as sterile as possible. First, the animals were anesthetized with a mixture of isoflurane (5 % for induction and 2 % for maintenance), and O_2_ (0.6 l/min) and N_2_O (0.4 l/min) using Harvard Apparatus equipment (Holliston, United States). A temperature probe soaked in vaseline was inserted rectally to control the body temperature between 36.6 - 37.5 °C. The eyes were covered with Bepanthen®(Bayer, Leverkusen, Germany) to prevent the cornea from drying out. After the surgeries, the mice received 10 % glucose (0.5 ml/30 g body weight, *s.c.*) as a fluid replacement, buprenorphine *i.p.* (0.01 μg/30 g body weight; Temgesic, Essex Pharma, Munich, Germany) and tramadol hydrochloride (Grünenthal GmbH, Aachen, Germany) to the drinking water (100 mg/200 ml) for three days, including the surgery day.

#### Cortical craniotomy

The cranial window procedure was performed as previously described with few modifications (Cupido et al., 2014). Briefly, after disinfecting the area with 70 % ethanol and removing the tissue, a 3-4 mm-diameter craniotomy was made over the somatosensory cortex (3.4 mm posterior to bregma and mediolateral 1.5 mm) using a driller. The drilling procedure was stopped at short intervals to remove bone particles and cool down the area with cortex buffer (in mM) 125 NaCl, 5 KCl, 10 glucose, 10 HEPES, 2 CaCl_2_, 2 MgSO_4_ [pH ~ 7.4] to prevent overheating of the underlying cortex. Next, using forceps (Fine Science Tools, Foster City, United States) the remaining bone was carefully removed to avoid damaging the meninges. Small bleeding was stopped with sponges (Gelastypt, Sanofi Aventis, Paris, France). A coverslip was placed on the exposed brain and the edge was sealed with dental cement (RelyX®, 3M ESPE, Saint Paul, United States). Finally, a custom-made metal holder (5 mm diameter) was put over the coverslip and glued with dental cement onto the bone.

#### Middle cerebral artery occlusion

Focal cerebral ischemia (FCI) was induced according to the Koizumi method of MCAO (Koizumi et al., 1986). In short, the left common carotid artery (CCA) and the external carotid artery were permanently ligated with silk sutures. A silicone-coated filament (Doccol Corp, Sharon, United States) was introduced through an arteriotomy and advanced into the right internal carotid artery until mild resistance was felt, indicating the filament reached the origin of the MCA to occlude the blood flow. After 15 min occlusion, the filament was gently withdrawn and a suture was made around the CCA, to prevent backflow through the arteriotomy. After recovery from anesthesia, the mice were kept in their cages with free access to food and water.

#### Neurological Score

Mice were evaluated for neurological deficits using the modified Bederson Score System (Bederson et al., 1986; Bieber et al., 2019). The animals were examined for motor impairments 24 h after the MCAO as listed below (Table 3; Fig. 1A). Animals presenting a score of 5 were euthanized and excluded from the experiment.

**Table 3.** Bederson neurological score (0-5)

| Score | Mouse behavior |
| --- | --- |
| 0 | no observable deficit |
| 1 | forelimb flexion |
| 2 | forelimb flexion and decreased resistance to lateral push |
| 3 | circling |
| 4 | circling and spinning around the cranial-caudal axis |
| 5 | no spontaneous movement |

### Immunohistochemical analysis

#### Preparation of vibratome slices

After experimental procedures, mice were deeply anesthetized with Ketamine (Ketavet®, Pfizer, New York City, United States) / Xylazine (Rumpon®, Bayer Healthcare, Leverkusen, Germany) in 0.9 % NaCl (100 μl/10 μg body weight; *i.p*.) and dissected by performing a bilateral axillary thoracotomy to expose the heart. A butterfly needle was inserted into the left ventricle and perfusion was started with 1x phosphate buffered saline (PBS) followed by 4 % paraformaldehyde (PFA) in phosphate buffer. Simultaneously, an incision of the superior vena cava allowed the blood to drain off. The brains were carefully removed and postfixed in the same fixative solution overnight at 4 °C.

The fixed brains were sliced in PBS into coronal sections (40 μm) at a Leica VT1000S vibratome (Leica Biosystems, Wetzlar, Germany). The coronal sections at the hippocampal level (Bregma −1.34 mm to −2.70 mm according to Franklin and Paxinos, 1997) were collected in 48-well culture plates containing PBS and used for free-floating immunohistochemistry.

#### Antibody incubation

Vibratome sections were incubated in blocking solution (0.3 % Triton X-100 and 5 % horse serum in PBS) for 1 h at room temperature (RT). Next, the sections were incubated with polyclonal rabbit anti-Iba1 antibody (1:1000, Wako Chemicals USA, Richmond, United States), diluted in blocking solution, overnight at 4 °C. On the next day, the sections were washed 3× for 10 min with PBS and incubated with the anti-rabbit secondary antibody (1:1000; Alexa Invitrogen, Walthan, United States), diluted in blocking solution, for 2 h at RT. Additionally, 4′,6-Diamidin-2-phenylindol (DAPI, *f.c*. 1 μg/ml, AppliChem, Darmstadt, Germany), a specific dye for nucleic acids, was added to the secondary antibody incubation. After washing with PBS, the slices were placed in a water bath and mounted with Immu-Mount™ (Thermo Fisher Scientific, Walthan, United States).

#### Fluoro-Jade C staining

To monitor MCAO-induced cell death, Fluoro-Jade C (FJC) staining was used, which is an established detection technique for degenerating neurons (Schmued and Hopkins, 2000). Vibratome sections, collected in 0.1 M PBS, were mounted on gelatin-coated slides and then dried on a slide warmer at 50 °C for 30 min. The slides containing the sections were immersed in a basic alcohol solution consisting of 1 % sodium hydroxide in 80 % ethanol for 5 min. Next, they were rinsed for 2 min in 70 % EtOH, for 2 min in distilled water, and then incubated in 0.06 % potassium permanganate solution for 15 min. Following a 1 min water rinse, the slides were transferred for 20 min to a 0.0001 % solution of FJC (Millipore, Burlington, United States) dissolved in 0.1 % acetic acid vehicle and kept in the dark. Thereafter, slides were washed in distilled water three times for 1 min, air dried, and coverslipped with DPX (Sigma-Aldrich, St. Louis, United States). FJC staining images were taken with the automated slide scanner microscope (Axio Scan.Z1, Zeiss, Jena, Germany).

### Microscopy

#### Automated epifluorescence microscopy on fixed brain slices

Epifluorescent images were taken by the automated slide scanner AxioScan.Z1 (Zeiss, Jena, Germany) equipped with an LED Light Source Colibri 7 (Zeiss, Jena, Germany). A Plan-Apochromat 10×/0.45 objective for pre-focusing and a Plan-Apochromat 20×/0.8 objective for fine focus image acquisition were applied with appropriate emission and excitation filters. Offline image stitching (8 μm stacks, variance projection) for overviews of brain slices and further analysis was performed using ZEN 1 Software Black Edition (Zeiss, Oberkochen, Germany).

#### Confocal laser-scanning microscopy

Confocal images were taken using a laser-scanning microscope (LSM-710, Zeiss, Jena, Germany) with a Plan-Apochromat 40×/1.4 Oil DIC (UV) VISIR M27 objective. Z-stacks of images taken at 1 μm intervals were processed with Fiji/ImageJ software (Schindelin et al., 2012) and displayed as maximum intensity projections for analyses with the MIA Software (provided by Prof. Bart Eggen, University of Groningen, Netherlands). The images were taken in the cortex, hippocampus, and thalamus in both brain hemispheres.

#### 2-Photon laser-scanning microscopy

High-resolution *in vivo* imaging was performed using a custom-made two photon laser-scanning microscope (2P-LSM) equipped with a mode-locked Ti:Sapphire laser (Vision II, Coherent, Santa Clara, United States). Scanning and image acquisition were controlled using custom-written software ScanImage (Pologruto et al., 2003). Additionally, the setup was equipped with XY-galvanometer-based scanning mirrors (Cambridge Technology). The excitation wavelength of the laser was set at 890 nm and a 20×/1.0 water-immersion objective (W Plan-Apochromat, Carl Zeiss, Jena, Germany) was used. To minimize photodamage, the average excitation laser intensity was kept at a minimum for a sufficient signal-to-noise ratio, ranging from 30 to 50 mW depending on depth. The emitted light was detected by photomultiplier tubes (H10770PB-40, Hamamatsu Photonics, Hamamatsu, Japan). The imaging settings were selected every time equally (256 × 256 px, Zoom 2, frame rate 1.9 Hz).

#### Microglia imaging *in vivo*

For microglia imaging *in vivo*, isoflurane anesthesia was used as described. After sedation, the animals were fixed with a metal holder on a custom-made head restrainer. Before the imaging session was started, animals were injected with circa 50 μl Texas Red-dextran (50 mg/ml, 70 kDa; Invitrogen, Waltham, United States), via tail vein injection, for visualization of brain blood vessels. The marked blood vessels served as a map to image the same region of interest (ROI) over several sessions. Time-lapse imaging of cortical microglia was performed by the repeated acquisition of fluorescence image stacks (15 focal planes with 2 μm axial spacing) recorded 50 - 70 μm below the dura mater. Subsequent image stacks were recorded every 60 s, totalizing 20 stacks acquisition from a single ROI. The mice were imaged before (baseline), 30 min, 24, and 48 h after the MCAO (Fig. 1B).

#### Ca^2+^ imaging *in vivo* in slightly anesthetized mice

First, the animals were anesthetized as described. After fixing the animal’s head, the isoflurane was reduced to 0.5 % until the end of the imaging session. Astrocytic GCaMP3 signals were recorded on a single focal plane located in the somatosensory cortex at a depth of 50 - 70 μm inferior to the dura mater. Three to four ROIs were recorded for each mouse. The same experimental design was used as described above (Fig. 1D). All images and movies were further processed using Fiji/ImageJ software.

#### Ca^2+^ imaging in acute brain slices

For acute brain slice imaging, the animals were sacrificed by decapitation 3 h after MCAO. First, the brain was dissected and placed in an ice-cooled, carbogen-saturated (5 % CO_2_ - 95 % O_2_, pH 7.4) cutting solution, containing (in mM) 87 NaCl, 3 KCl, 1.25 Na_2_H_2_PO_4_, 25 NaHCO_3_, 25 Glucose, 75 Sucrose, 3 MgCl_2_, and 0.5 CaCl_2_. Next, the brain tissue was coronally cut (300 μm slice thickness) using a vibratome (Leica VT 1200S, Wetzlar, Germany) and transferred into a nylon basket as a slice holder with incubation solution, containing (in mM) 12.6 NaCl, 3 KCl, 1.2 Na_2_H_2_PO_4_, 25 NaHCO_3_, 15 Glucose, 2 MgCl_2_, and 1 CaCl_2_ 35 °C for 30 min. No more than 5 slices were collected from each brain. After incubation, the slices were let to recover with continuous oxygenation for at least 30 min at RT before recording. Subsequently, slices were transferred to an imaging chamber under the 2P-LSM and kept submerged by a custom-made platinum grid with nylon threads for mechanical stabilization. The imaging chamber was continuously perfused with oxygenated perfusion solution, containing (in mM) 12.6 NaCl, 3 KCl, 1.2 Na_2_H_2_PO_4_, 25 NaHCO_3_, 15 Glucose, 2 MgCl_2_, and 1 CaCl_2_ at a flow rate of 2-5 ml/min. The imaging sessions were done in both hemispheres (contra- and ipsilateral) and the chosen ROIs were located in the somatosensory cortex at a depth of 50 μm. For the investigation of the CBD effect over time, the slice incubated in oxygenated perfusion solution was imaged at 0, 10, 20 and 30 min. At the end of the period of 30 min the perfusion solution was switched to perfusion solution containing CBD 100 μM and the ROI was recorded again using the same schedule described above (Fig. 1E). Afterwards each application, the slice was removed and the chamber was washed completely with perfusion solution. CBD was first dissolved in a stock solution of 10 mM in dimethyl sulfoxide (DMSO), which was then added to the perfusion solution to obtain the desired 100 μM final concentration. CBD concentration was based on previous studies (Castillo et al., 2010).

### Data analysis

#### Staining data

FJC staining was analyzed, first, via measurement of fluorescence intensity on the total area of the brain slice, to obtain the percentage (%) of affected area per slice (Fig. 2C). Next, the number of FJC-positive cells was counted in the hippocampal dentate gyrus (GD) and CA1 area, thalamus and striatum (Fig. 2D, F, G). The ROIs were determined by drawing the respective structures on the slice. All slices were analyzed at the same coronal level and using the same ROI position and size.

**Figure 2.**
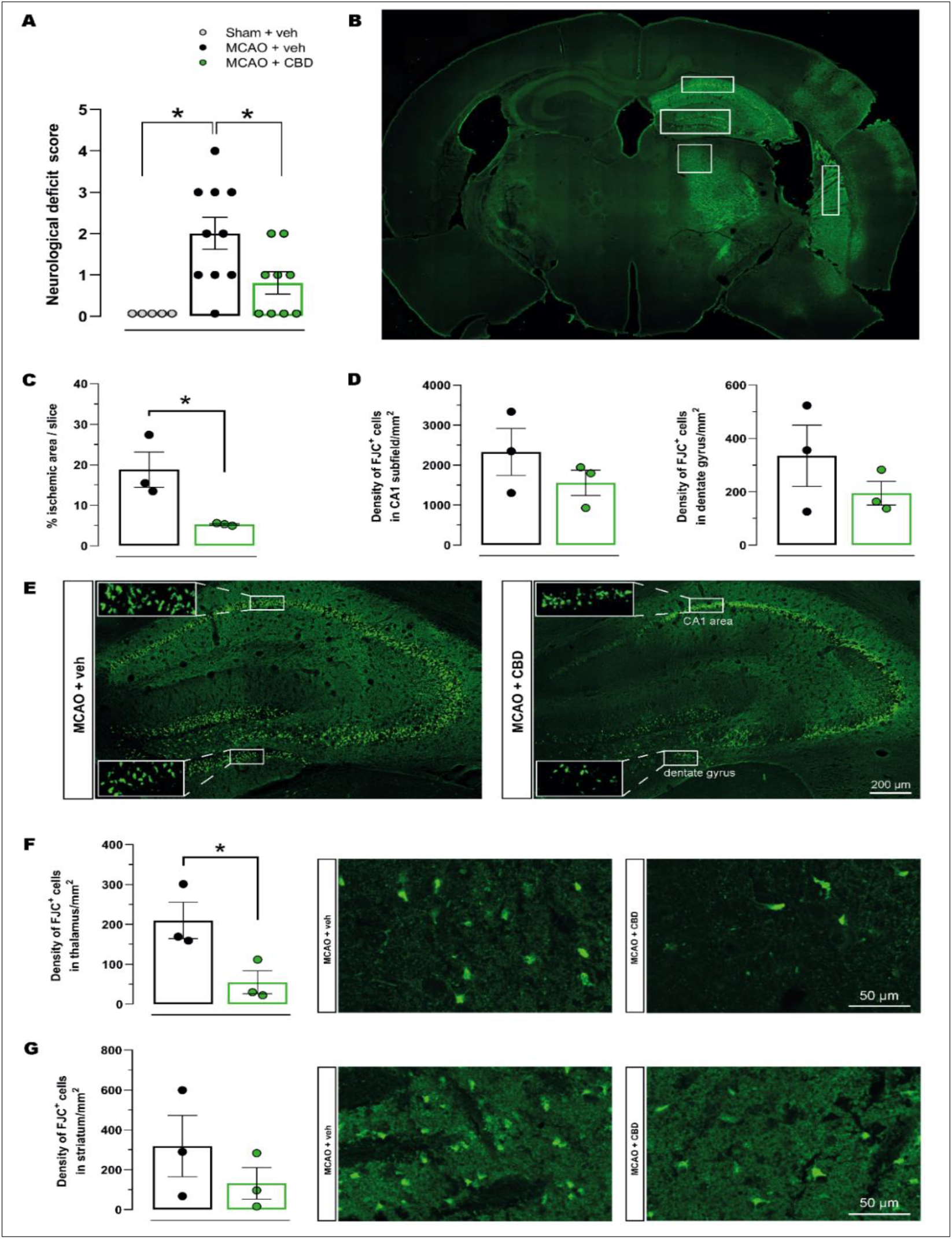
Cannabidiol protected against neurological impairments and neuronal death induced by MCAO. **A** Neurological impairment was evaluated 24 h post-ischemia using the neurological deficit score. **B** Representative figure illustrating a coronal brain section at the intermediate level of the hippocampus showing the CA1 subfield, DG, thalamus, and striatum where the analysis for FJC was performed. **C** Percentage (%) of FJC-positive neurons per slice. **D** The density of FJC-positive neurons in the hippocampal CA1 area and DG. **E** Representative photomicrographs of FJC-positive cells in the CA1 and DG in the different experimental groups. **F** Density of FJC-positive neurons in the thalamus and respective representative photomicrographs. **G** The density of FJC-positive neurons in the striatum and respective representative photomicrographs among the groups. Stats: data are shown as individual values (dots) and the means ± SEM (columns and bars) of the experimental groups (*n*= 3-10/group). \**p* ≤ 0.05. DG: dentate gyrus; FJC: Fluoro-Jade C; MCAO: middle cerebral artery occlusion.

#### MIA Software

MIA is a semi-automatic morphological parameter extractor for single-microglia images. Sholl profiles detailing about 23 parameters of microglia morphology can be obtained after analyses with MIA. This software was developed by Prof. Bart Eggen and his team (University of Groningen, Groningen, Netherlands). The cortex, hippocampus and thalamus in both brain hemispheres (contra- and ipsilateral side) were investigated. For each mouse, three microglia cells per brain region were analyzed.

#### Analyses of Ca^2+^ imaging data

Ca^2+^ imaging data were analyzed, using the custom-made MATLAB application *MSparkles* (Stopper et al., in preparation). MSparkles was specifically designed for the visualization and analysis of cellular events, visualized using fluorescence indicator dyes. A novel algorithm to estimate fluorescence fluctuations at basal concentrations of signaling molecules (*F*_0_), such as Ca^2+^ or Na^+^, removes fluorescence background and enables the detection of microdomain events. *F*_0_ was computed by fitting a polynomial curve along the temporal axes of each pixel. Before polynomial fitting, statistically large values were removed. Subsequent event detection and analysis were based on the range projection of the normalized and detrended image stack (Δ*F*/*F*_0_). Fluorescence events were detected as temporally correlated, local brightness peaks, and segmented into individual ROIs. ROI traces were integrated by computing the mean fluorescence (Δ*F*/*F*_0_) per ROI per recorded time point. These ROI traces were then subjected to peak analysis for transient extraction and classification, based on a transient’s peak amplitude.

Before analysis, Ca^2+^ imaging data were run through a pre-processing pipeline performing de-noising using the PURE-LET algorithm, image registration as well as a temporal median filter of size 3.

#### Statistical analyses

Prism 8 software (GraphPad, San Diego, CA, USA) was used for the statistical analysis. Data were examined for assumptions of a normal distribution using the Shapiro-Wilk normality test. In case of a normal distribution, the data were analyzed using Student’s *t*-test and one- or two-way analysis of variance (ANOVA) as appropriate, followed by the Tukey multiple-comparison post hoc test. In the two-way repeated measures ANOVA, the group was the between-subject factor, and time (test day) was the within-subject factor. For data sets with a non-normal distribution, the non-parametric Wilcoxon matched-pairs signed rank and Mann-Whitney tests were applied, as appropriate. For calcium-signals data, outlier detection with GraphPad Prisms Rout method (Q = 1 %) was performed. The data are expressed as mean ± SEM or median with interquartile range (IQR). Values of *p* ≤ 0.05 were considered statistically significant.

## RESULTS

### Ischemia-induced neurodegeneration and neurological deficits are reduced after cannabidiol treatment

The effects of ischemia and CBD on neurological function were evaluated 24 h after reperfusion by the neurological deficit score, as shown in Fig. 2A. The one-way ANOVA revealed significant differences in the scores among groups (*F*_2,21_ = 7.92, *p* < 0.05). Tukey’s post hoc analyses revealed that the sham+vehicle group had a lower score compared with the MCAO+vehicle group (*p* < 0.05). The MCAO+CBD group exhibited a significantly lower score compared with the MCAO+vehicle group (*p* < 0.05).

Neurodegeneration was evaluated by the increase in the number of dying neurons assessed by FJC staining two days after sham or MCAO surgery in the hippocampal CA1 subfield, DG, thalamus, and striatum, as illustrated in Fig. 2B. An increased percentage of FJC-positive cells per slice was found in the MCAO+vehicle compared with the MCAO+CBD group (Student’s *t*-test, *t*_4_ = 3.10, *p* < 0.05), reflecting the beneficial effect of CBD on MCAO-induced neurodegeneration (Fig. 2C). As shown in Fig. 2D, there were no statistical differences in FJC staining of hippocampal CA1 area and DG between the groups (Student’s *t*-test, *t*_4_ = 1.13 - 1.16, *p* > 0.05). However, a significant loss of neurons in the MCAO+vehicle group was detected when compared to the MCAO+CBD group (Student’s *t*-test, *t*_4_ = 2.87, *p* < 0.05; Fig. 2F). There was no statistical difference in the FJC-positive cell number in the striatum between the groups (Student’s *t*-test, *t*_4_ = 2.87, *p* > 0.05; Fig. 12G).

### Reduction of ischemia-caused microglia activation by cannabidiol treatment

MCAO-induced microglia activation was assessed by analyzing Iba-1 immunoreactivity in the cortex, hippocampus and thalamus comparing contra- and ipsilateral sides from animals’ brains (Fig. 3A). As shown in Fig. 3C, ischemia did not affect the soma area, total branch length, and number of nodes of microglia in the cortex (*F*_2,9_ = 1.08 - 2.45, *p* > 0.05) and thalamus (*F*_2,9_ = 5.85 - 10.85, *p* > 0.05) from the contralateral side of the lesion. However, the one-way ANOVA revealed significant differences in microglial soma area, total branch length, and the number of nodes in the hippocampus from the contralateral side among groups (*F*_2,9_ = 5.53 - 27.62, *p* < 0.05). Further analysis revealed that these three parameters were increased in the MCAO+vehicle group when compared with the sham+vehicle group (*p* < 0.05). Cannabidiol reversed this ischemia-induced effect in the hippocampus compared with the MCAO+vehicle group (*p* < 0.05).

**Figure 3.**
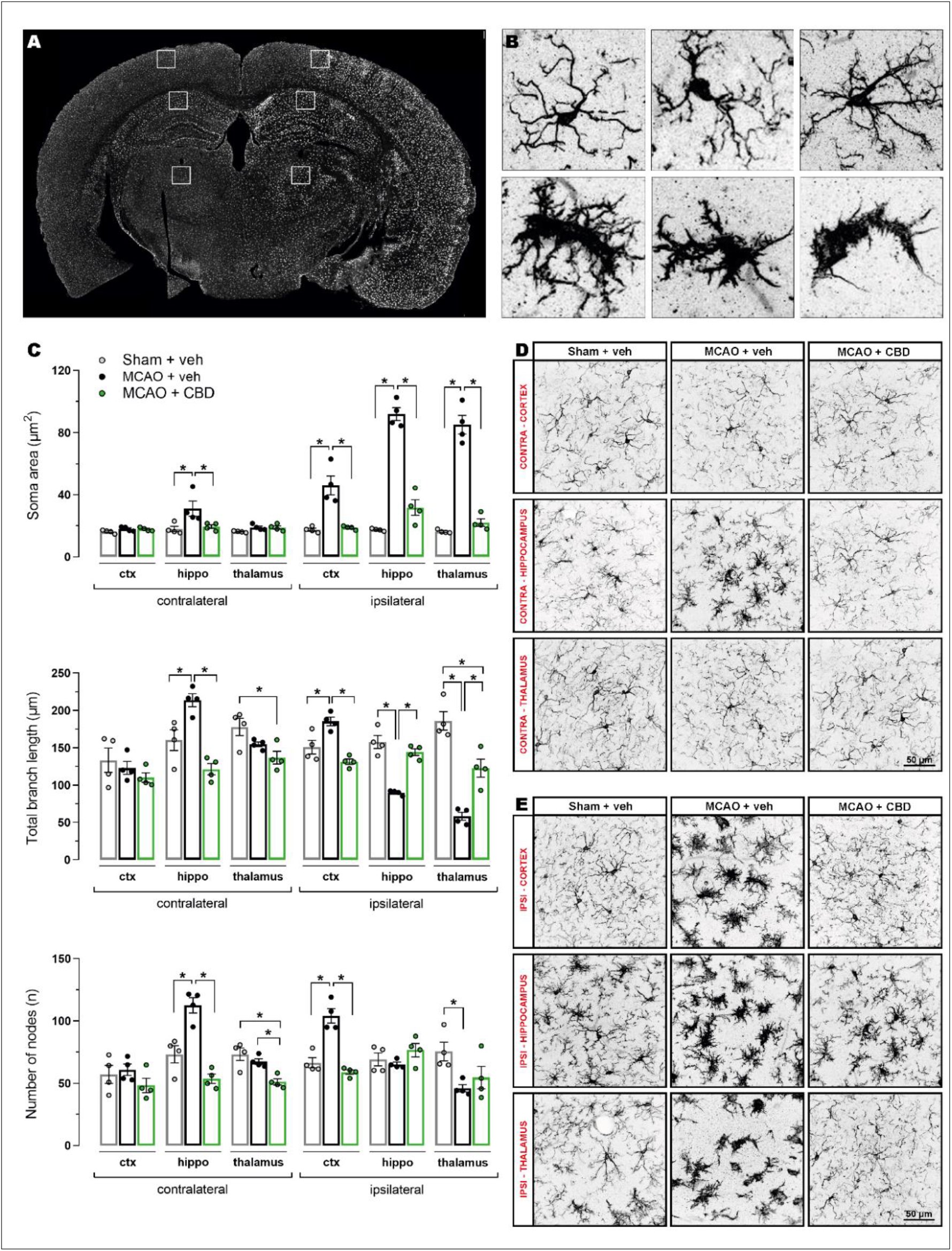
Cannabidiol reduced microglial activation induced by MCAO. **A** Representative figure illustrating a coronal brain section showing the cortex, hippocampal CA1 subfield and thalamus where the analysis for microglial morphology was performed. **B** Heterogeneity in the morphology of microglial cells was found among the groups. **C** Soma area, total branch length and number of nodes in the cortex, hippocampus and thalamus on the contra- and ipsilateral side among the groups. **D** Representative photomicrographs of Iba-1-IR cells on the contralateral side among the experimental groups. **E** Representative photomicrographs of Iba-1-IR cells on the ipsilateral side among the experimental groups. Stats: data are shown as individual values (dots) and the means ± SEM (columns and bars) of the experimental groups (*n*= 4/group). \**p* ≤ 0.05. MCAO: middle cerebral artery occlusion.

Ischemia-induced microglial activation was also observed on the ipsilateral side of the brain (Fig. 3C). The one-way ANOVA revealed significant differences in the soma area of microglia in the cortex, hippocampus, and thalamus among groups (*F*_2,9_ = 104.0 - 111.2, *p* < 0.05). Tukey’s post hoc test showed that the MCAO+vehicle group exhibited an increase in these parameters in all three brain regions compared with the sham+vehicle group (*p* < 0.05). CBD treatment alleviated this effect compared with the MCAO+vehicle group (*p* < 0.05).

Significant statistical differences were also found in the total branch length in the cortex, hippocampus, and thalamus on the ipsilateral side among groups (*F*_2,9_ = 16.97 - 39.41, *p* < 0.05; Fig. 3C). A significant increase in the total branch length in the cortex was found in the MCAO+vehicle group compared with the sham+vehicle group (*p* < 0.05). In the hippocampus and thalamus, the opposite ischemia-induced effect was observed, comparing the MCAO+vehicle with sham+vehicle group (*p* < 0.05). Ischemic mice that were treated with CBD exhibited a significant decrease in the total branch length in the cortex, and an increase in this parameter in the hippocampus and thalamus compared with the MCAO+vehicle group (*p* < 0.05). The one-way ANOVA also revealed significant differences in the number of nodes in the cortex and thalamus of the ipsilateral side of the lesion among groups (*F*_2,9_ = 4.72 - 33.14, *p* < 0.05). Tukey’s post hoc test showed that the MCAO+vehicle group exhibited an increase in the number of nodes in the cortex and a decrease in the thalamus compared with the sham+vehicle group (*p* < 0.05). CBD treatment alleviated the ischemia-induced increase in the number of nodes of microglia in the cortex compared with the MCAO+vehicle group (*p* < 0.05).

### I*n vivo* 2P-LSM demonstrates a reduced microglial activation in ischemic animals treated with cannabidiol

The microglial reaction during *in vivo* 2P-LSM recording in each group is shown in Fig. 4B. The two-way ANOVA revealed a main effect of group for the number of cells: *F*_2, 5_ = 6.60, *p* < 0.05) and soma area (*F*_2,5_ = 6.81, *p* < 0.05). A main effect of time (*F*_3,15_ = 7.65, *p* < 0.05) and a significant group × time interaction (*F*_6,15_ = 8.01, *p* < 0.05) was found for the soma area. At baseline conditions, no between-group differences were found for the number of cells and soma area, indicating no microglial activation before the injury for all experimental groups (Tukey’s post hoc test, *p* > 0.05). Comparing the MCAO+vehicle with sham+vehicle group at 30’ after MCAO, the ischemic animals presented more cells and increased soma area (Tukey’s post hoc test, *p* < 0.05). The ischemia-induced microglial activation was prevented by CBD treatment. The longitudinal analysis indicated that both the number of cells and the soma area significantly decreased at 30’ after MCAO in the MCAO+CDB group compared with the MCAO+vehicle group (*p* < 0.05). Similar outcomes for the MCAO+CBD *vs*. MCAO+vehicle groups were found for both parameters at 24 h after MCAO (*p* < 0.05). In the sham-operated and ischemic treated with CBD groups, the number of cells and soma area did not differ between the recorded time points, indicating no microglial activation along time (*p* > 0.05). However, the MCAO+vehicle group presented more cells and increased microglial soma area at 30’ and 24 h after MCAO compared to baseline conditions (*p* < 0.05).

**Figure 4.**
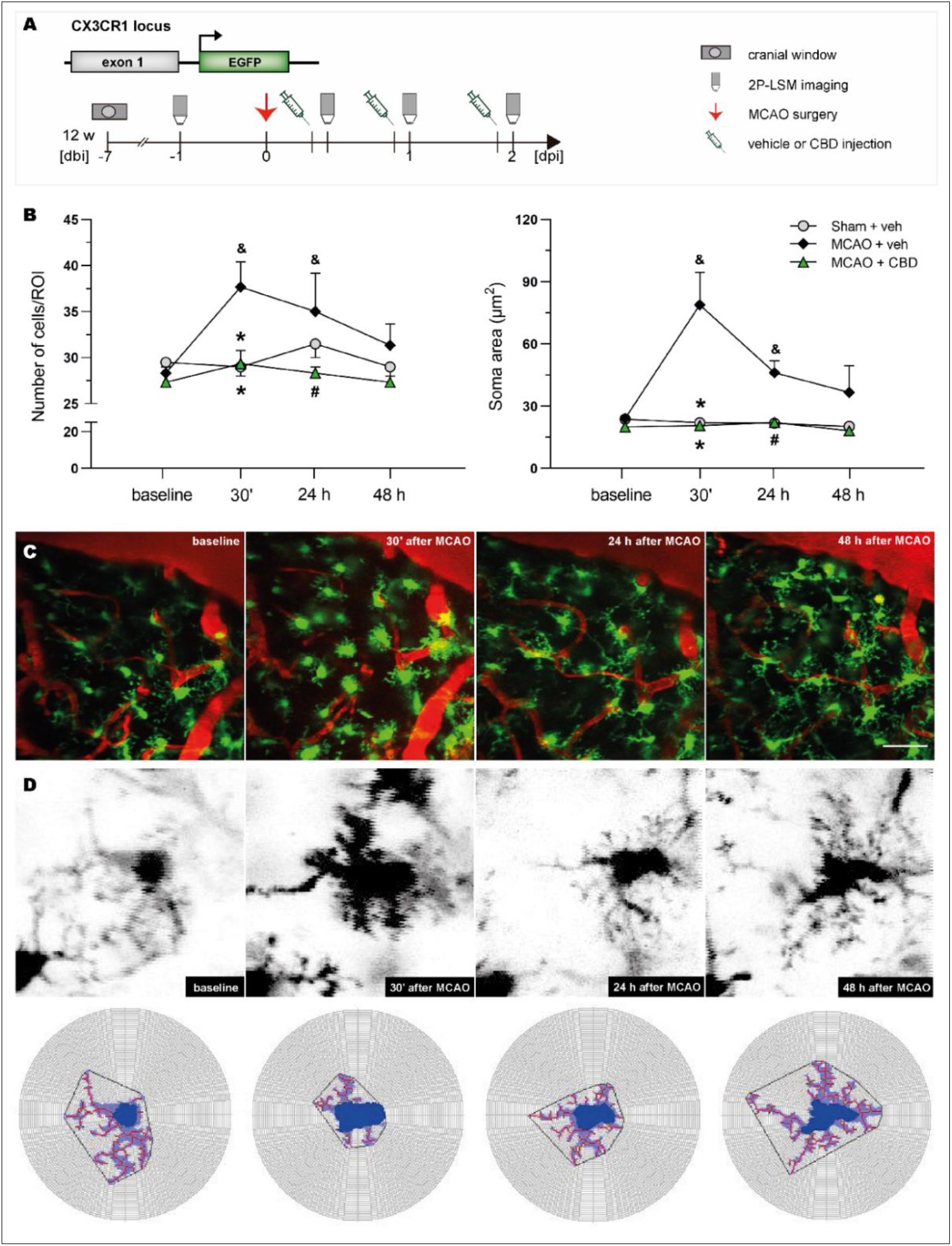
Cannabidiol prevented microglia activation in the somatosensory cortex of MCAO animals. **A** *In vivo* microglia imaging was performed after vehicle or cannabidiol injection during baseline, 30 min, 24, and 48 h after reperfusion. **B** Temporal distribution of number and soma area of microglia in the somatosensory cortex on the ipsilateral side among the groups. **C** Representative maximum-intensity projection of one stack of one field of view at recorded time points from MCAO + veh group. **D** Representative photomicrographs of microglia from an ischemic mouse at recorded time points and respective Sholl analysis with MIA software. Stats: the bars represent the means ± SEM of the experimental groups (*n*= 3/group). \**p* ≤ 0.05 compared to MCAO + veh group at 30’, ^#^*p* ≤ 0.05 compared to MCAO + veh group at 24 h and ^&^*p* ≤ 0.05 compared to MCAO + veh group at baseline. Scale bar = 50 μm. MCAO: middle cerebral artery occlusion.

### Cannabidiol treatment did not change astrocytic calcium signaling *in vivo* after ischemia

Astrocytic calcium signaling was evaluated *in vivo* by visualization of the genetically encoded Ca^2+^ indicator GCaMP3. This experiment aimed, first, to determine whether MCAO changes the amplitude and duration of astrocytic Ca^2+^ signals. Secondly, we examined whether treatment with CBD modulates Ca^2+^ signals in the somatosensory cortex on the ipsilateral side of the brain of ischemic mice.

As shown in Fig. 5A and B, neither amplitude nor duration of Ca^2+^ signals was changed over the days analyzed in the group treated with vehicle (within group comparison, *W* = −6, *W* = −6 – −4, *p* > 0.05, respectively). Similarly, no difference in the signal amplitude (within group comparison, *W* = −6 – 0, *p* > 0.05) and duration (within group comparison, *W* = −6 – −4, *p* > 0.05) was found over time in the group of ischemic mice treated with CBD. Comparing the signals obtained in the MCAO+vehicle group with those obtained in the MCAO+CBD group, in each time point, no difference in the signal amplitude and duration were found (*U* = 1 – 4, *U* = 2 – 4, *p* > 0.05, respectively).

**Figure 5.**
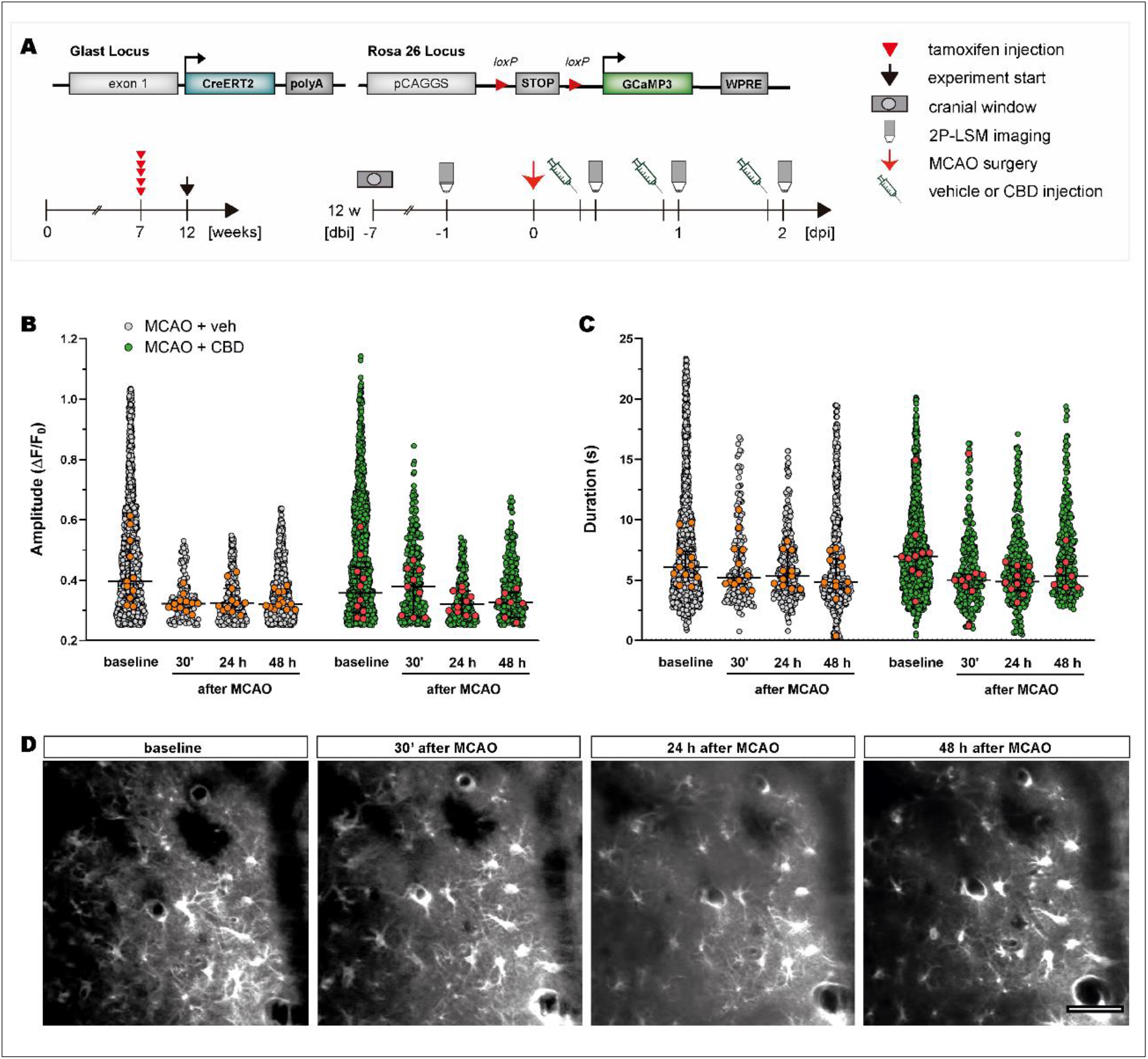
Cannabidiol did not modulate astrocytic Ca^2+^ signaling in the somatosensory cortex after MCAO. **A** GCaMP3 expression in astrocytes was induced five weeks before the experiment started. *In vivo* two-photon Ca^2+^ imaging was performed after vehicle or CBD injection during baseline, 30 min, 24, and 48 h after MCAO. **B-C** Amplitude and duration of astrocytic Ca^2+^ signals on the ipsilateral side of the brain after MCAO among groups. **D** Representative single frames of *in vivo* two-photon recordings of one FOV at recorded time points. Scale bars = 50 μm. Stats: data points represent FOV of the experimental groups (*n* = 3/group) and are displayed as median ± IQR. Grey and green data points in the background display single Ca^2+^ signals. CBD: cannabidiol; FOV: field of view; MCAO: middle cerebral artery occlusion.

### Astrocytic calcium signals are not affected in the early phase of focal ischemia

To confirm the results obtained in the experiment with GLAST^GCaMP3^ mice *in vivo*, we performed imaging of Ca^2+^ calcium signals *ex vivo*. Firstly, Ca^2+^ signals in the somatosensory cortex from both brain hemispheres were compared. Contra and ipsilateral side did not display differences in the amplitude (*U* = 12, *p* > 0.05) and duration (*U* = 15, *p* > 0.05) of astrocytic Ca^2+^ signals (Fig. 6B).

**Figure 6.**
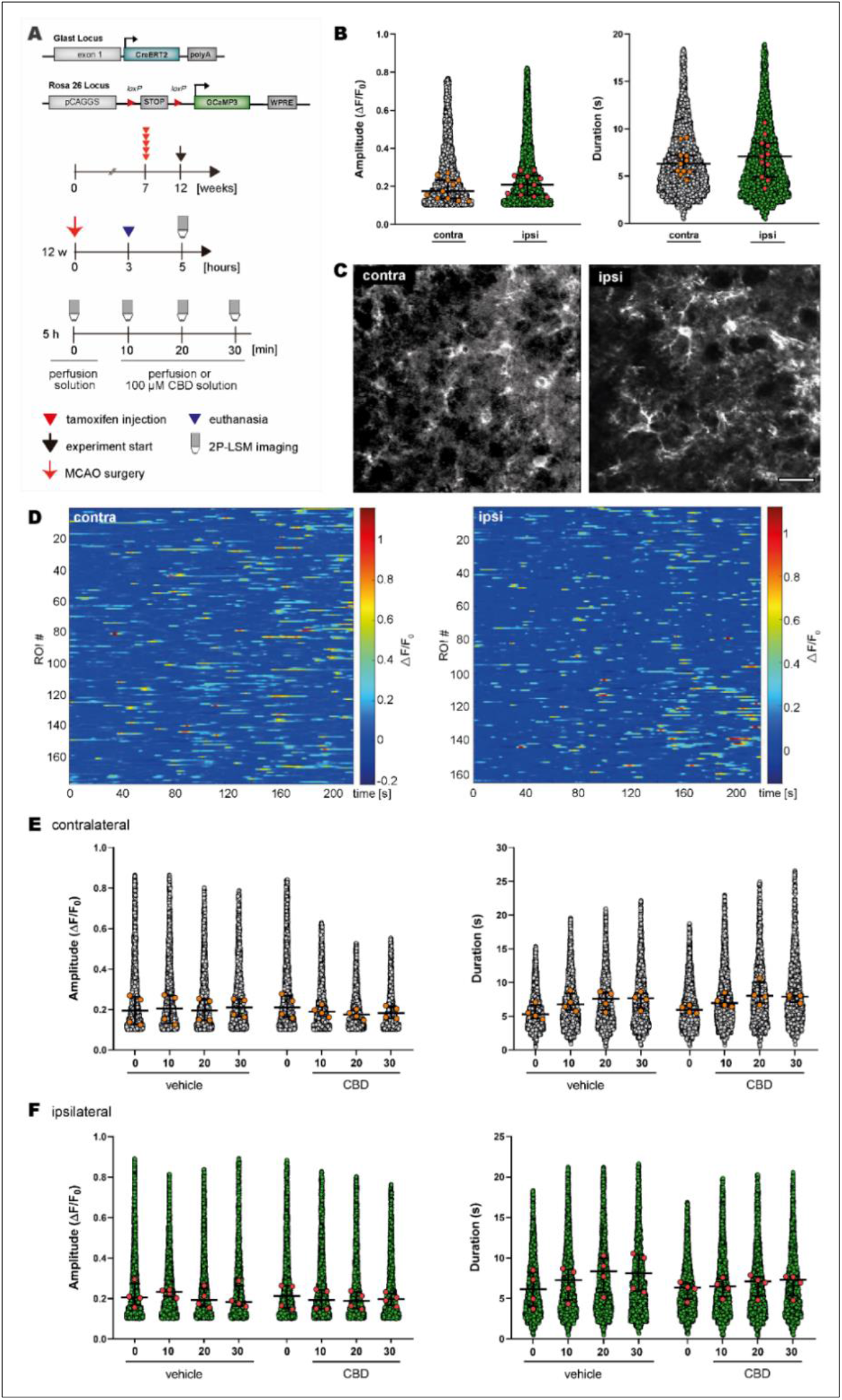
Cannabidiol did not change astrocytic Ca^2+^ signaling in the somatosensory cortex of acutely isolated slices of ischemic mice. **A** GCaMP3 expression in astrocytes was induced five weeks before the experiment started. *Ex vivo* two-photon Ca^2+^ imaging was performed 5 h after MCAO at different time points in the presence of perfusion or CBD solution. **B** Amplitude and duration of Ca^2+^ signals in astrocytes on the brain’s contra- and ipsilateral side after MCAO. **C** Representative single frames of ex *vivo* two-photon recordings of one FOV on the contra- and ipsilateral side. **D** Representative heatmaps of fluorescence amplitude (ΔF/F_0_) from the contra- and ipsilateral side of the brain. **E-F** Amplitude and duration of astrocytic Ca^2+^ signals over time in the presence of perfusion or CBD solution on the brain’s contra- and ipsilateral side, respectively. Stats: data points represent FOV of the experimental groups (*n*= 4-5/group) and are displayed as median ± IQR. Grey and green data points in the background display single Ca^2+^ signals. Scale bar = 25 μm. CBD: cannabidiol; FOV: field of view; MCAO: middle cerebral artery occlusion.

To investigate if Ca^2+^ signals could be modulated by the presence of CBD, the acutely isolated brain slices were incubated, firstly, with perfusion solution and then CBD solution. As shown in Fig. 6E, on the contralateral side, the amplitude and duration of Ca^2+^ signals were not changed after incubation with perfusion solution comparing the time points analyzed (within group comparison, *W* = 2 – 4, *W* = 10, *p* > 0.05, respectively). Similarly, no difference in the signal amplitude and duration was found when the slices were incubated with CBD solution (within group comparison, *W* = −8 – −6, *W* = 8 – 10, *p* > 0.05, respectively). Finally, no difference in the signal amplitude and duration was found over time comparing the signals obtained after incubation with perfusion solution and CBD solution (*W* = −6 – 8, *W* = 4 – 8, *p* > 0.05, respectively).

Fig. 6F shows that, on the ipsilateral side, there was no change in the Ca^2+^ signal’s amplitude over time in the presence of perfusion solution (within group comparison, *W* = −2 – 2, *p* > 0.05) or CBD solution (within group comparison, *W* = −8 – −4, *p* > 0.05). Similarly, no significant difference was found for the duration of signal overtime when the slices were incubated with perfusion solution (within group comparison, *W* = 10, *p* > 0.05). Comparing the signal duration after incubation with CBD solution no difference was found among the time points recorded (within group comparison, *W* = 4 - 10, *p* > 0.05). No significant difference in the amplitude nor duration of Ca^2+^ signals was found comparing the incubation with perfusion solution and CBD solution (*W* = −8 – 0, *W* = −6 – −2, *p* > 0.05, respectively).

## DISCUSSION

In the present study, we found that CBD (10 mg/kg, *i.p.* 30 min, 24, and 48 h after ischemia) reverted neurological deficits that was induced by FCI in mice. CBD treatment reduced by 75 % the ischemia-induced neurodegeneration area. CBD also reduced neuroinflammation in ischemic mice, reflected by *in situ* and in *vivo* decreases in reactive microglia.

Impairment of sensorimotor and cognitive performance is a common outcome in rodents with MCAO (Hattori et al., 2000; Truong et al., 2012; Linden et al., 2014). In the present study, the neurological impairments were detected in mice in the subacute phase of injury, *i.e.*, one day after MCAO, using the Bederson score system. CBD attenuated the effects of MCAO, reflected by a decrease in the neurological score, indicating an improvement in neurological function. Consistent with our results, CBD led to an improvement of motor and neurological deficits in mice submitted to 4 h MCAO and treated immediately before and 3 h after the occlusion. These effects of CBD were accompanied by a reduction in infarct size and an increase in cerebral blood flow (Hayakawa et al., 2004; Mishima et al., 2005; Hayakawa et al., 2007). Another study demonstrated that CBD (5 mg/kg, *i.p.*) given once 15 minutes after reperfusion led to the functional and sensorimotor recovery in neonatal rats submitted to MCAO (Ceprián et al., 2017). Similar beneficial effects of CBD on neurological function were also observed in MCAO rats treated with CBD, *i.c.v.*, for 5 days before surgery (Khaksar and Bigdeli, 2017). On the other hand, mice submitted to 4 h MCAO and treated with CBD (3 mg/kg) from day 5 did not show improved neurological score and motor coordination on day 14 after reperfusion. These results indicate that the therapeutic time window is an important feature of CBD treatment in CI, with neuroprotective actions occurring in the subacute phase of ischemia.

MCAO is known to cause a robust reduction in cerebral blood flow and consequent massive neuronal death. Accordingly, we found significant neurodegeneration (detected by FJC staining) in the hippocampus, thalamus and striatum of MCAO animals, which paralleled neurological impairments in those animals. The short-term CBD treatment reduced the extension of neuronal loss induced by MCAO, especially in the thalamus. Other studies have shown reduced neuronal death in the striatum and hippocampus after CBD treatment in mice (Hayakawa et al., 2008) and gerbils (Braida et al., 2003) submitted to focal and global CI, respectively. Moreover, a reduction of necrotic neurons in the cortex was observed in newborn pigs submitted to a hypoxic-ischemic brain injury and treated with CBD (1 mg/kg, *i.v*.) 30 min after the ischemic insult (Pazos et al., 2013). Notably, CBD (10 mg/kg, *i.p*. 30 min before and 3, 24, and 48 h after ischemia) decreased neurodegeneration and normalized caspase-9 protein levels 21 days after GCI in mice, demonstrating that the neuroprotective action of CBD led to sustained beneficial effects after the injury (Mori et al., 2017).

Neuroinflammation is a critical aspect of stroke, which includes the early activation of microglia and production of cytokines and chemokines (Iadecola and Anrather, 2011; Kim *et al.*, 2016; Jayaraj et al., 2019). Depending on injury severity, microglia may present distinct functional and spatiotemporal-dependent profiles in CI, which may protect or contribute to the ischemic injury evolution (Yasuda et al., 2011; Benakis et al., 2015; Fumagalli et al., 2015). We have observed an extensive microglial activation 2 days after MCAO, which extended from the ipsilateral side of the brain, including large areas in the cortex, hippocampus, striatum, and thalamus, to the contralateral hippocampus. In line with our results, activated microglia were abundant in the cortex and thalamus 2 days after MCAO in rats. In the same study, early microglial activation was also observed in regions outside of the MCA territory, such as the contralateral cortex and hippocampus (Morioka et al., 1993). In this sense, microglial reactivity not only indicates imminent ischemic neuronal damage but possibly may reflect subtle changes in neuronal activity outside the MCA territory. Selective neuronal loss, which refers to the death of single neurons with preserved extracellular matrix, *i.e.*, in the ischemic penumbra zone, is consistently associated with microglial activation in the first few days after injury (Yasuda et al., 2011; Emmrich et al., 2015; Park et al., 2018). As both these processes, *i.e.*, selective neuronal death and activation of microglia, affect the salvageable penumbra, hindering functional recovery after reperfusion, they represent potential therapeutic targets in ischemic stroke (Hughes et al., 2010; Baron et al., 2014).

CBD mediated-neuroprotection after experimental CI has generally been related to the modulation of inflammation, including the control of microglia activation, and the toxicity exerted by these cells by producing pro-inflammatory mediators (Pazos et al. 2013; Mohammed et al., 2017; Mori et al., 2017). In mice submitted to 4 h MCAO, repeated CBD treatment from day 1 after ischemia reduced the number of Iba1-positive cells expressing HMGB1 a proinflammatory cytokine that is massively released during the acute phase of ischemic processes (Hayakawa et al., 2009). A reduction in the number of Iba1-positive cells was also reported in neonatal rats submitted to a model of ischemic stroke and treated once with CBD (5 mg/kg, *i.p*.) 15 min after the injury (Ceprián et al., 2017). Our results also demonstrated the anti-inflammatory potential of CBD, as shown by a potent reduction in microglial activation in several cerebral areas 2 days after the onset of ischemia.

To test whether the effects of CBD on microglia could be also detected *in vivo*, we performed time-lapse imaging of microglial activity using the 2P-LSM. Soon after reperfusion, we have observed that microglia become activated and increased in number in the ischemic penumbra. Supporting our findings, *in vivo* imaging of mice submitted to a cortical microhemorrhage has shown a coordinated pattern of microglia migration, where microglia within 200 μm of the injury migrated toward the lesion, leading to an increased microglia local density (Ahn et al., 2018). Moreover, in mice submitted to MCAO, it was demonstrated that many round-shaped microglia migrated to the peri-infarct area 24 h after the insult (Tanaka et al., 2003). According to our results and also using an *in vivo* approach in mice, Jolivel and coworkers (2015) have shown that all microglia in the penumbra were found associated with blood vessels within 24 h post reperfusion. The authors have also demonstrated that these perivascular microglia started to phagocytose endothelial cells, leading to an activation of the local endothelium and contributing to the degradation of blood vessels with an eventual breakdown of the BBB. Considering these findings, the inhibition of microglial activation within the first day after stroke could stabilize blood vessels in the penumbra, improving the outcomes after ischemia through an increase in the blood flow (Jolivel et al., 2015). Our results show that CBD treatment decreased microglia activation after MCAO, which is consistent with other findings and reinforces the strong anti-inflammatory profile of CBD in ischemic conditions.

Besides microglia, astrocytes play an active role in the neuroinflammation process as well, producing complex and not yet completely understood responses after CI (Liu and Chopp, 2016). In newborn piglets submitted to hypoxia-ischemia, treatment with CBD attenuated the loss of cortical GFAP-positive cells and decreased the levels of S100β in the cerebrospinal fluid (Lafuente et al., 2011). In mice submitted to GCI, CBD (10 mg/kg, *i.p*.) decreased the hippocampal reactivity of astrocytes (GFAP-positive) and total levels of GFAP 21 days after the insult (Mori et al., 2017). In cultured astrocytes, CBD treatment decreased the ß-amyloid-induced release of proinflammatory mediators such as nitric oxide, TNF-α, S100B, and IL-1β. In the same study, CBD treatment (10 mg/kg, *i.p*.) for 15 days diminished the pro-inflammatory response and gliosis triggered by the intrahippocampal injection of ß-amyloid (Esposito et al., 2011).

We tested whether focal ischemia would impact astrocytic Ca^2+^ signaling, a characteristic form of excitability in this cell type, and whether CBD treatment could modulate these signals. Unexpectedly, ischemic mice did not display significant differences in cortical astrocytic Ca^2+^ signals up to 2 days after MCAO. Additionally, no difference in astrocytic Ca^2+^ signals was observed in the somatosensory cortex between contra- and ipsilateral brain sides. Our results are not in line with others showing an increase in astrocytic Ca^2+^signals after CI. For example, in mice submitted to photothrombotic-induced FCI for 20 min, *in vivo* imaging showed an increase in frequency and amplitude of transient Ca^2+^ signals in astrocytes. The authors reported, an increased astrocytic Ca^2+^ signal in the penumbra although it was smaller than that in the core region (Ding et al., 2009). Moreover, permanent MCAO led to an increased astrocytic Ca^2+^ activity in the penumbra of aged mice, while a reduction of astrocytic Ca^2+^ activity was observed in adult mice (Fordsmann et al., 2019). A strong increase of intracellular Ca^2+^ in astrocytes associated with detrimental peri-infarct depolarizations was also reported by Rakers and Petzold (2017) in mice after permanent MCAO. The reason we did not detect a significant difference in astrocytic Ca^2+^ signals after MCAO is unknown. Considering that neurons are more sensitive to ischemia than astrocytes, it is possible that the short occlusion time applied in our study was insufficient to elicit astrocytic responses in the somatosensory cortex after the ischemia. Yet, the cited studies have examined astrocytic Ca^2+^ signaling *in vivo* using a permanent model of FCI or photothrombotic-induced FCI, which display significant differences with the transient FCI model used in this study, mostly for not allowing cerebral reperfusion. Our results involving the absence of astrocytic activation in the somatosensory cortex after MCAO remain elusive and factors other than those mentioned above could be implicated in the absence of astrocytic response found in our study.

One limitation of the present study was that we did not distinguish between resident brain microglia and eventually infiltrating peripheral macrophages. Whether treatment with CBD might differently impact the microglial subpopulations after MCAO remains to be determined.

Overall, the present findings suggest that the functional and structural protective effects of CBD are closely associated with anti-inflammatory action in the subacute phase of ischemia.

